# Cell Type Composition Analysis: Comparison of statistical methods

**DOI:** 10.1101/2022.02.04.479123

**Authors:** Sean Simmons

## Abstract

Measuring changes in cell type composition between conditions (disease vs not, knockout vs wild type, treated vs not, etc) is fast becoming a standard step in single cell RNA-Seq analysis. Despite that, there is no agreement on the best approach for this type of analysis. As such, we decided to test numerous methods for cell type composition analysis, seeing how they performed in terms of false positive rate and power. Though there is not one clear winner, we do find two method (the propeller method with asin normalization and Dirichlet regression with the alternative parameterization) perform well in most situations. Most importantly, consistent with results in differential expression analysis, we see that it is important to take into account sample to sample (mouse to mouse, person to person, etc) variability to avoid high false positive rates. We also see evidence that aggregation (aka pseudobulk) based method slightly outperform the mixed model methods we tested.

## Introduction

When analyzing single cell and nuclei data, one is often interested in testing if the cell type composition changes between two (or more) conditions (also known as differential abundance testing)—that is to say we want to see if two conditions have the same cellular make up. The question of compositionality comes up in many fields outside single cell genomics (microbiome [1], etc), and there are many methods for such analysis, ranging from those in stats 101 (Fisher exact and chi squared) to those built on more complicated statistical models (dirichlet regression, multinomial mixed models, etc) and many in between.

We aimed to test a large swath of these methods to see if they have nice properties for single cell analysis. In particular, we wanted to test false positive and true positive rates, if the p-values are correctly distributed, run time, and robustness to down sampling.

### Properties of the different methods

Each method has different properties, some of which are important in some cases, some of which are important in other cases. Below are a few of those properties we are most interested in:

#### Modelling of sample to sample variability

It is well known that there are sample to sample (animal to animal and cell line to cell line, for example) differences in single cell RNA-seq data [2], both due to biological differences (slightly different ages, differences in genotype, slight differences in environments, randomness during development, etc) and technical ones (slightly differences in dissection and dissociation, differences in quality of data between 10X experiments, etc). Many methods ignore this sample to sample variability, but others take it into account.

Note the methods that correct for sample to sample variability tend to break into 2 types: 1) aggregation (aka pseudobulk) based methods, where the cells in each sample are reduced to one measurement (usually the number or percent of cells in a given cell type for that sample) and analysis is run on these per sample statistics or 2) mixed model methods, where random terms specific to each sample are added into the cell specific analysis. It is worth noting that a similar divide occurs in single cell differential expression methods [2].

#### Modelling of compositionality

For single cell data, since one is only sampling a subset of cells in a given region, the total number of cells in each cell type isn’t usually meaningful, only the percent of cells in each cell type is (since even within samples of the same type there is likely to be large variability in the number of cells recovered). Since these percentages have to add up to 100%, however, that means if one cell type goes up in composition others must go down. This can lead to false positive results if it is not taken care of—for example, if one cell type disappears then all other cell types increase in proportion. It is worth noting that, in some cases, it might be acceptable to include these ‘false positives’ (they are actually changing in proportion, they are just not likely to be the most interesting change biologically) but in other cases it might not be. One way to work around this issue is by using a base cell type (see below).

#### Requirement of base cell type

Some methods require the choice of one cell type as a base cell type, and changes in all other cell types are measured relative to that one. There are some plus sides to this (it indirectly helps to correct for effects of compositionality, easy to interpret, etc), but in many cases it can be hard to choose a particular cell type as the base, something we discus more in the conclusion.

#### Correcting for confounders

Some methods make it easy to correct for confounders (by including them as an extra covariate, etc), some do not. This can be important in many situations, such as in human data where one often has individuals of different ages, sexes, etc, in the same analysis.

### Description of methods

We decided to implement many methods, most (but not all) of which have been used in single cell analysis before. Each method was given a short abbreviated name, and the methods are described below (in no particular order, with the abbreviated name in bold). Citations are included for those that we are aware of having been used in single cell analysis before (likely to have missed some) and for particular implementations we used.

#### Poisson

This a Poisson regression based method [3]. For each cell type, the number of cells in each sample is calculated then a Poisson regression is run with an offset term equal to the log of the total number of cells in the corresponding samples. We used the glm function in R.

##### NB

NB is a negative binomial regression based method. It is very similar to the Poisson method, except negative binomial regression is used instead of Poisson. We used the glm.nb function in the MASS package [4].

##### Chi

The basic chi-squared test, run on a different 2 by 2 table for each cell type with one dimension being condition, the other indicating if a given cell is the cell type of interest or not. We used the chisq.test function in R.

##### Fisher

Same as Chi, except with the Fisher exact test instead of the chi squared [5]. We used the fisher.test function in R.

#### Logistic

For each cell type run a logistic regression (a slight modification of https://www.nxn.se/valent/2020/11/28/s9jjv32ogiplagwx8xrkjk532p7k28), where the dependent variable is 1 if the given cell is the cell type of interest, 0 otherwise. P-values are extracted from the glm function in R.

#### Logistic_mixed

Same as logistic, except with a random term corresponding to sample of origin, similar to the MASC method [6]. We used the glmer function in lme4 [7].

#### Dirichlet

Runs a Dirichlet regression with the standard parameterization, where the dependent variables are the sample by sample cell type proportions for each cell type [5]. We used the DirichReg function in DirichletReg [8].

#### Dirich_alt

Same as Dirichlet except using the alternative parameterization, which requires a choice of base cell type. We used the DirichReg function in DirichletReg [8].

#### Wilcox_Percent

For a given cell type, for each sample we calculate the percent of cells in that sample that come from the given cell type, then use a Wilcox test to compare these between conditions [9]. This was run with the wilcox.test function in R.

#### Wilcox_Base

Same as Wilcox_Percent, except instead of dividing cell number by total number of cells to get percentages, we divide through by the number of cells in a particular (base) cell type. This was run with the wilcox.test function in R.

#### Multinomial

Fits a multinomial model with cell type as the output [10]. Used the mblogit function in the mclogit package [11].

#### Multinomial_mixed

Same as Multinomial, except with a random variable corresponding to sample of origin. We used the mblogit function in the mclogit package [11].

#### propel_logit/propel_asin

Based off the propeller method implemented in the speckle package (https://github.com/Oshlack/speckle) [12], these methods use a linear model of transformed proportions to estimate effects. propel_logit uses a logit transformation, propel_asin uses an asin transformation. Note we slightly modified the code to run more smoothly with our pipeline, but mathematically it should produce the same results.

#### Multi_overdisp_v1, Multi_overdisp_v2, Multi_overdisp_v3, Multi_overdisp_v4, Multinomial_mixed_overdisp

Same as Multinomial, except with over dispersion (a different method for each one), except for Multinomial_mixed_overdisp which is a mixed version (note that this actually is identical to Multinomial_mixed and will be removed later). Note this is only included in the overdispersion section. We used the mblogit function in the mclogit package.

Note we left out many methods here. There are numerous slight variations on the above (such as replacing the wilcox test with a t-test or another test in Wilcox_Percent approach, or using DESeq/EdgeR/etc instead of the standard NB test) we did not include due to limited bandwidth, but which may make a large difference in the analysis. There are also some more recent single cell methods, namely scDC [13], which involve reclustering using the transcriptional information. Though this seems promising, we left it out and focused only on methods that did not require access to the underlying transcriptional information (since our pipeline did not support it). There are also Bayesian methods (scCODA [14], etc) which, though of great interest, we did not include since our analysis focused on more frequentist concepts (p-values, FDR, power, etc). We also did not consider methods that dealt with soft clustering (so approaches built for methods like topic modelling where each cell can belong to multiple topics/clusters). Finally, there are some very recent methods that attempt to find differences in cell type composition without having labelled clusters ahead of time (such as Milo [15]). These are an exciting new direction, but our work focuses on the more traditional approach (if anything in single cell analysis RNA-seq analysis can be considered traditional) of finding differences in composition of known cell type clusters (moreover, these new methods are currently being compared independently of our work: https://github.com/singlecellopenproblems/SingleCellOpenProblems/issues/140)

Table 1 indicates, for each method, what properties it has. Note that, though it may not explicitly be taken into account, methods with base cell types do indirectly deal with compositionality. In general, it is preferable in most cases to have methods with the properties (so those with lots of Y’s) as opposed to those without them (those with lots of N’s).

**Table 1:** List of properties for tested methods

| Method | Models compositionality explicitly? | Models sample to sample variability? | Doesn't require base variable? | Easily corrects confounders? |
| --- | --- | --- | --- | --- |
| Logistic | N | N | Y | Y |
| Chi | N | N | Y | N |
| Fisher | N | N | Y | N |
| Dirichlet | Y | Y | Y | Y |
| Wilcox_Percent | N | Y | Y | N |
| Wilcox_base | N | Y | N | N |
| logistic_mixed | N | Y | Y | Y |
| Poisson | N | Y | Y | Y |
| multinomial | Y | N | N | Y |
| multinomial_mixed | Y | Y | N | Y |
| propel_logit | N | Y | Y | Y |
| Dirich_alt | Y | Y | N | Y |
| NB | N | Y | Y | Y |
| propel_asin | N | Y | Y | Y |

We can see that only one method (dirichlet) has all the desired properties, while many (multinomial_mixed, propel_logit, propel_asin, dirich_alt, nb, poisson, logistic mixed, and wilcox_percent) have 3 out of 4 (some of these require a base cell type, others do not correct for compositionality).

### Testing for false positives and p-values

Our first test is to see what happens when there is no signal in our data—in theory we should get uniformly distributed p-values. To do this, we take some single cell datasets consisting of control samples (so samples that are not known to have a disease, knockout, etc). In particular, we use:

1. Immune cell data: We took immune cell data from a recent paper on bacterial sepsis [16] and selected all sample that are controls and CD45 sorted. This left samples from 19 patients with a total of 36716 cells. For cell type, we used the Cell_Type metadata column in the data. For methods that require a base cell type we use the T cells.
2. Autism control data: We downloaded a single nuclei ASD dataset and selected all control samples that are > 11 years old from the PFC [17]. This left us with 8 samples and 26838 cells. For cell type, we used their cell type annotation and aggregated into 6 more general categories (Astroglia, Endo, Excitatory, Inhibitory, Microglia, and ODC). For methods that require a base cell type we use the Inhibitory neurons.

For each of these samples, we randomly assign half the samples to condition 1, the other half to condition 2 (if there are an odd number of samples condition 1 gets the extra sample) and use each method to test for cell type composition differences between the two conditions. We repeat this 1000 times (except for in our overdispersion analysis where we only ran it 100 times, see below), each time giving a different permutation of the condition labels.

Since the condition labels are random and the samples are all control/normal samples (note could still be confounders such as age, etc, in the data which we are ignoring but could have an effect on the results) then a good method should assign p-values as a uniform distribution on the range [0,1] (ignoring hidden confounders or outliers in the data, and the fact that there are relatively few permutations of the ASD data (8 choose 4 or 70 permutations)). To look at this, for each method we take the p-values calculated in each iteration for each cell type and plot the resulting distribution as a boxplot, see Fig 1a. If the distribution is uniform, we should expect a distribution where the median is around .5, the quantiles are around .25 and .75, and the ends of the whiskers are around 0 and 1. The closer the distribution moves to 0, the more false positives we will get—we say such a distribution is overinflated. Looking at the results, we see that methods that do not correct for sample to sample variability have distributions that are closer to 0, as does the Poisson method. All other methods, however, have pretty reasonable distributions in both datasets, with the wilcox based methods being particularly conservative in the ASD sample (aka prone to false negatives).

**Fig 1:**
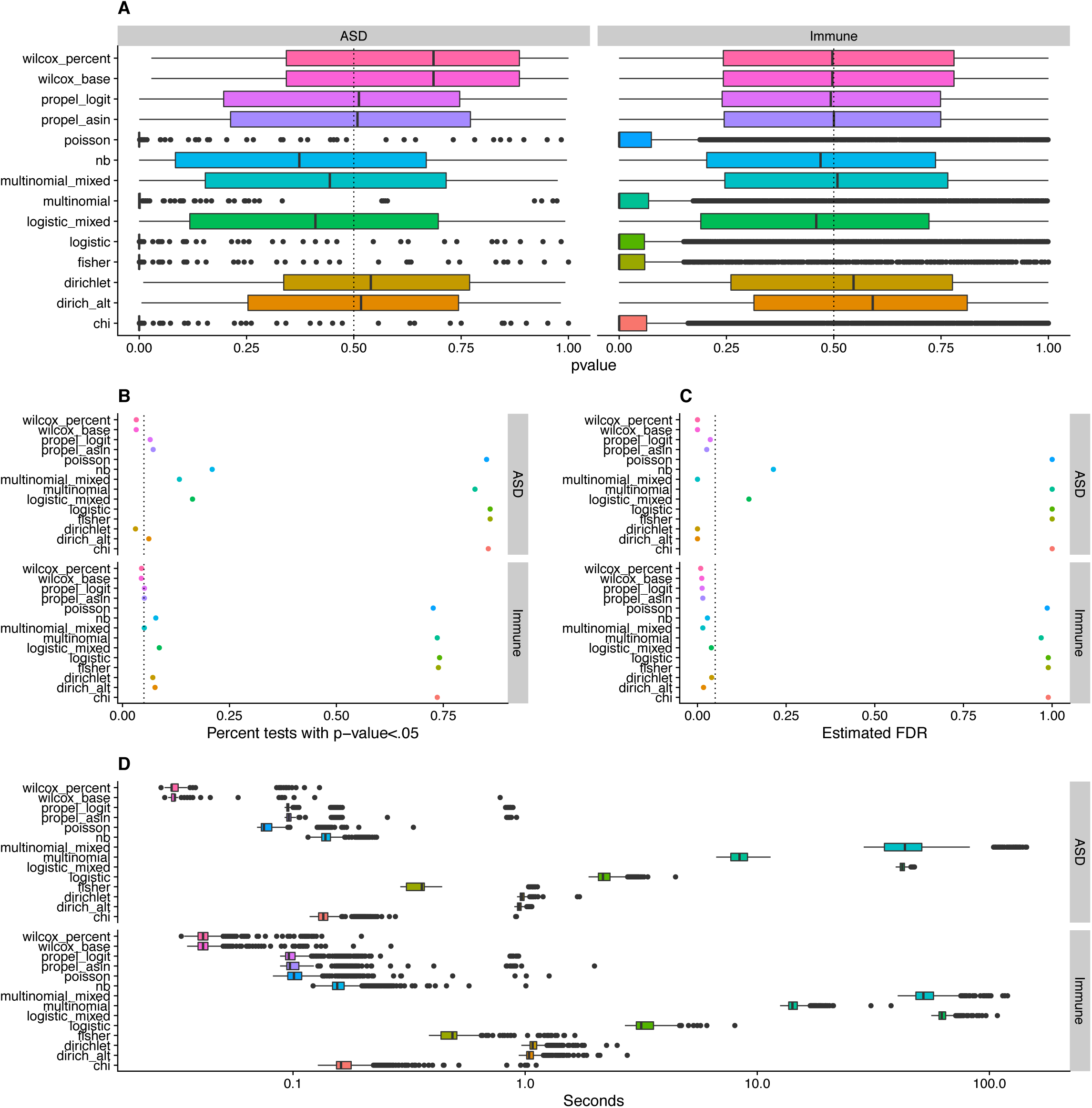
Method performance in the absence of cell type differences a) For each method, we collected the p-values of each celltype in each permutation and plotted the results as a box plot, with p-values on the x axis and methods on the y axis. On the left are the results for the ASD dataset, on the right the results for the Immune cell atlas. A well calibrated method should have it’s median around .5, it’s quantiles around .25 and .75, and it’s ends around 0 and 1. We see that Poisson regression and methods that do not take into account sample to sample variability have smaller than expected p-values, while the remaining methods perform well. b) For each method we calculate the percent of tests with uncorrected p-value<.05 for both the ASD dataset (top) and Immune atlas dataset (bottom). A well calibrated method should be around .05. We see that Poisson regression and methods that do not take into account sample to sample variability have much higher percentages than expected. c) We estimated the real FDR for each method from applying an FDR cutoff of .05. A well calibrated method should have FDR less than or equal to .05. We see that Poisson regression and methods that do not take into account sample to sample variability have higher than expected FDR values, while the remaining methods perform reasonably well in both datasets. d) For each method (on the y axis), we plot a box plot of runtimes over all 100 iterations in both the ASD dataset (top) and Immune data (bottom). The x axis is the runtime in seconds and log scaled. We can see a large variation between methods, with wilcox based methods being the fastest and mixed model methods being the slowest, though all methods run in <1 min on average.

We can also look at the percent of tests that reach the (nominal) .05 p-value cutoff—if the p-values are correct, about 5% of tests should have p-values <.05. The results of this are in Fig 1b. We see in the ASD datasets, the dirichlet, propel, and Wilcox based methods have the closest to 5% of their p-values being less than .05, while for the immune dataset the mixed multinomial, NB, and logisitic mixed model based methods also do well. Again, those methods that don’t take into account sample to sample variability do the worse, with Poisson also performing poorly.

Finally, we can look at the FDR—if we perform FDR correction and label all celltypes with FDR correct p-value <.05 as significant, how many hits will we get? If the p-values are doing well this number should be roughly .05, if not less (depending if the FDR correction method, BH correction in this case, is overly conservative or not). This is plotted in Fig 1c. We see inflation of the FDR in methods that do not account for sample to sample variability and poisson regression, though it seems to be reasonably controlled in both datasets by the wilcox based methods, the dirichlet based methods, the propel based methods, and the mixed multinomial based methods. The NB and logistic mixed model do well in the Immune dataset, but are slightly inflated in the ASD one.

We run a similar test, except we now include a randomly generated binary covariate in the analysis (for those methods that allow covariate correction, the others are run without correction). We see very similar results to Fig 1, except surprisingly the dirichlet method is suddenly over inflated in the ASD data, with an FDR close to .5, see Fig 2a-c.

**Fig 2:**
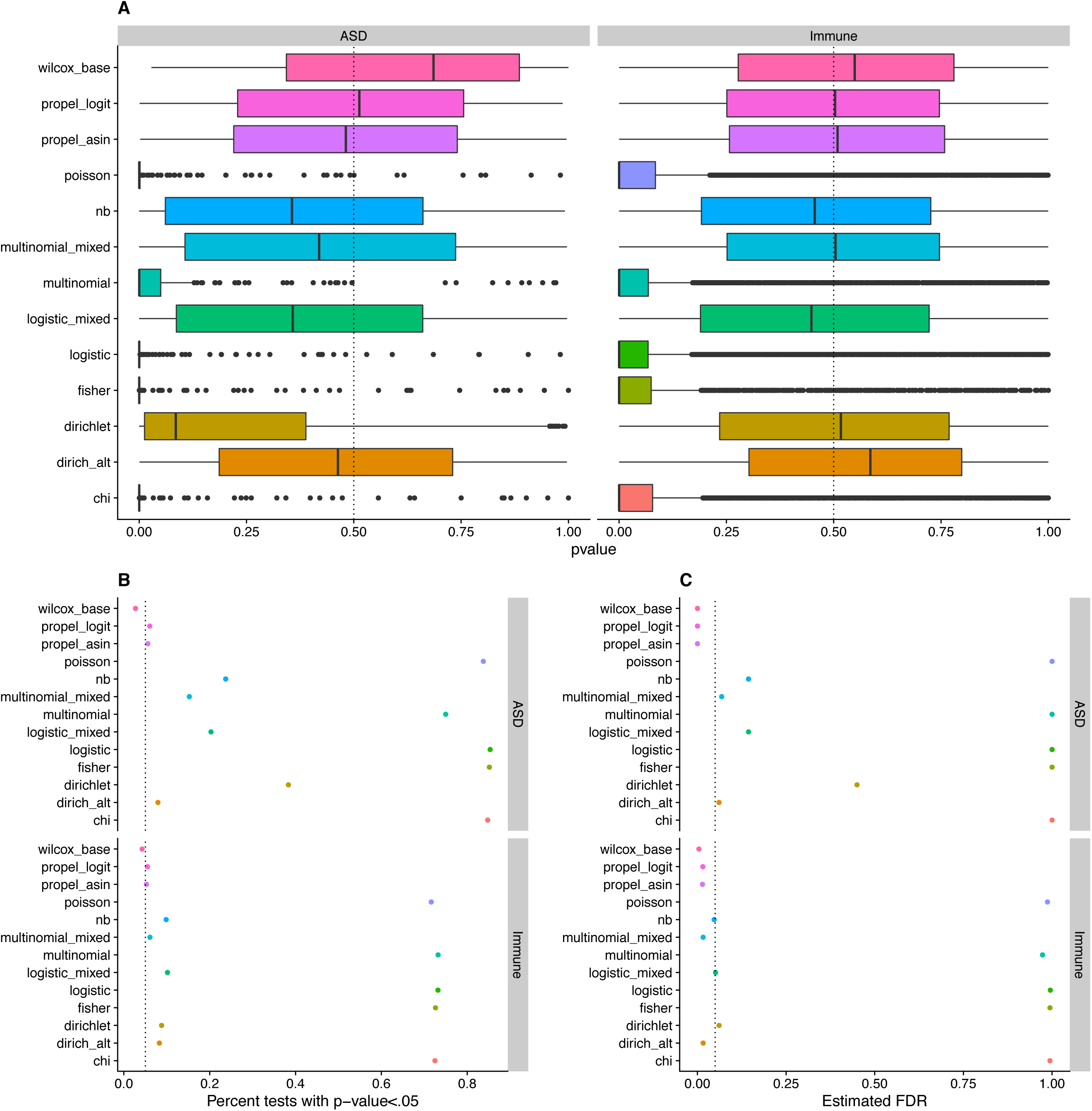
Method performance with correction for covariate a) For each method, we collected the p-values of each celltype in each permutation and plotted the results as a box plot after correcting for covariates, with p-values on the x axis and methods on the y axis. On the left are the results for the ASD dataset, on the righ the results for the Immune cell atlas. A well calibrated method should have it’s median around .5, it’s quantiles are .25 and .75, and it’s ends around 0 and 1. The results are similar to Fig 1a, with slightly lower p-value in the ASD dataset, with particularly large inflation of the dirichlet results. b) For each method calculate the percent on tests with p-value<.05 for both the ASD dataset (top) and Immune atlas datset (bottom) after correcting for a covariate. A well calibrated method should be around .05. The results are similar to Fig 1b, except slightly more inflation in the ASD dataset, particularly for the dirichlet. c) We estimated FDR for each method from applying an FDR cutoff of .05 after coring for a covariate. A well calibrated method should have FDR less than or equal to .05. The results are similar to Fig 1c, except slight inflation of FDR in the ASD data, particularly for dirichlet regression with an FDR near 50%.

### Runtime

One important consideration for these methods is the runtime. For each of the above experiments, we recorded the run time. In Fig 1d you can see the runtime for each method over all 1000 iterations of both datasets (note the log scale of the x axis). We can see a very large difference in runtime, with the longest being the mixed models at around a minute, and the shortest being the Wilcox based methods with runtimes in the realm of .01-.1 seconds, with the rest of the methods in between. Having said that, 1 minute runtimes on 19 samples and >36,000 cells is reasonable for many real world use cases, so all of these methods seem to be fast enough to be useable in many (arguably most) current applications. It is worth noting that these runtimes are based off one particular implementation of each method with one particular setting of parameters and includes preprocessing time—changing these could change the relative runtimes, but the above analysis is likely to give us some idea of how the runtimes compare.

### Power

One other important, but much harder to test, consideration is that of power—if there is a signal how likely is each method to detect it. Note this is also important since correcting for compositionality is likely to be most important in cases where there is a cell type composition change (unlike the conditions tested above).

To test this we used the same approach as above, except introduced a simulated change in cell type composition. We decided to use the ASD dataset, and simulated 2 changes in cell type—one where 50% of the excitatory neurons in condition 1 were removed at random, the other where 5% were removed at random. We then ran the same analysis as before. In theory we would like the Excitatory to have low p-values, the rest to have uniform p-values (note it is possible this might not be the desired behavior for all applications, but we believe it to be the one that makes the most sense in general). This is what we looked at in Fig 3, a-c. Note Fig 3a is stratified by % of excitatory neurons removed (.05 vs .5) and by cell type (True corresponding to cell types with a true change, False to the remaining cell types (non-Excitatory neurons)). Looking at the non-Excitatory cell types (the ones labelled False) we do see inflation of p-values for nb and logistic_mixed with the 50% change (and to a lesser extent 5%), which is unsurprising as these 2 methods do not correct for compositionality, but otherwise the results are similar the Fig 1. Note there is also an even smaller inflation of p-values in the propel based methods with the 50% change, thought it doesn’t seem to effect the false discovery rate for either propel_asin or propel_logit. We also tested the power of each method in Fig 3d. Here, power is the percent of iterations where the excitatory neurons are significant at an FDR of .05 (so where the true effect is detected). We can see that with a 5% change we have very low power among the methods that performed well on the FDR test, but with a 50% change we get much more power. Interestingly, the Dirichlet based methods have less power than the nb, propel, mixed model based, and even wilcoxen based methods in this particular test.

**Fig 3:**
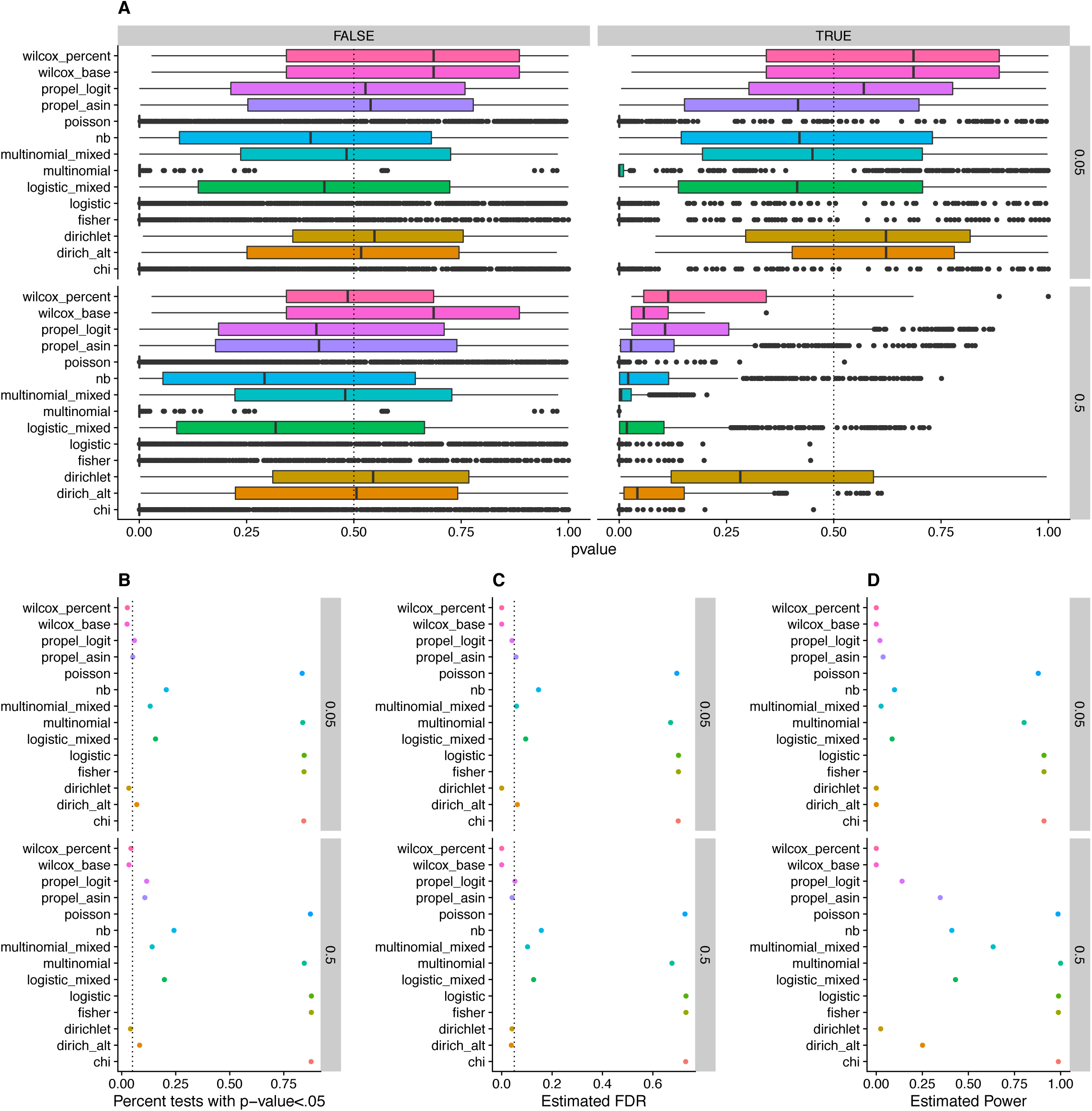
Method power to detect changes a) For each method, we collected the p-values of each celltype in each permutation and plotted the results as a box plot, with p-values on the x axis and methods on the y axis. The plot on the left correspond to cell types with no changes, those on the right to those with actual changes (aka excitatory neurons). The top of each plot corresponds to a small change (5% change), the bottom to a much larger one (50% change). A well calibrated method should have it’s median around .5, it’s quantiles are .25 and .75, and it’s ends around 0 and 1 for the cells with no change, but lower p-values for the cells with a signal. We see the results are similar to those in Fig 1a for the cells with no signal, except a moderate inflation of nb and logistic_mixed methods in the 50% dataset. b) For each method calculate the percent on tests (restricted to non-Excitatory cell types) with p-value<.05 for both the 5% dataset (top) and 50% dataset (bottom). A well calibrated method should be around .05. The results are similar to Fig 1b, except slight inflation of p-values in the 50% dataset, particularly for nb and logistic_mixed. c) Estimated FDR for each method from applying an FDR cutoff of .05 after coring for a covariate. A well calibrated method should have FDR less than or equal to .05. The results are similar to Fig 1c d) Estimated power for each method (percent of iterations where the FDR for excitatory neurons is <.05). Can see that there is more power to detect changes when the changes are bigger. In the 50% dataset we see a large variation between methods, with multinomial mixed model based methods outperforming other methods (ignoring methods with inflated FDR levels), particularly relative to the dirichlet based methods.

### Overdispersion vs sample to sample variability

One property we did not explore in the above was overdispersion—does accounting for overdispersion improve performance, even in the case with no sample to sample variability correction. Indeed, sample to sample variability can lead to a model that appears overdispersed. We did see in the above that nb outperforms poisson, where nb can be viewed as an over dispersed version of poisson (note: the “overdispersed” mutinomial mixed model is actually just the same as the mutinomial mixed model (the overdispersion flag is ignored)). To look more into this, we compared standard multinomial regression to 4 different overdispersed versions of multinomial regression. We ran it through the same pipeline as in Fig 1. The results are plotted in Fig 4. The main takeaway is that the overdispersion doesn’t seem to help in this case—the v4 overdispersion is actually worse than no overdispersion, while the rest are almost identical.

**Fig 4:**
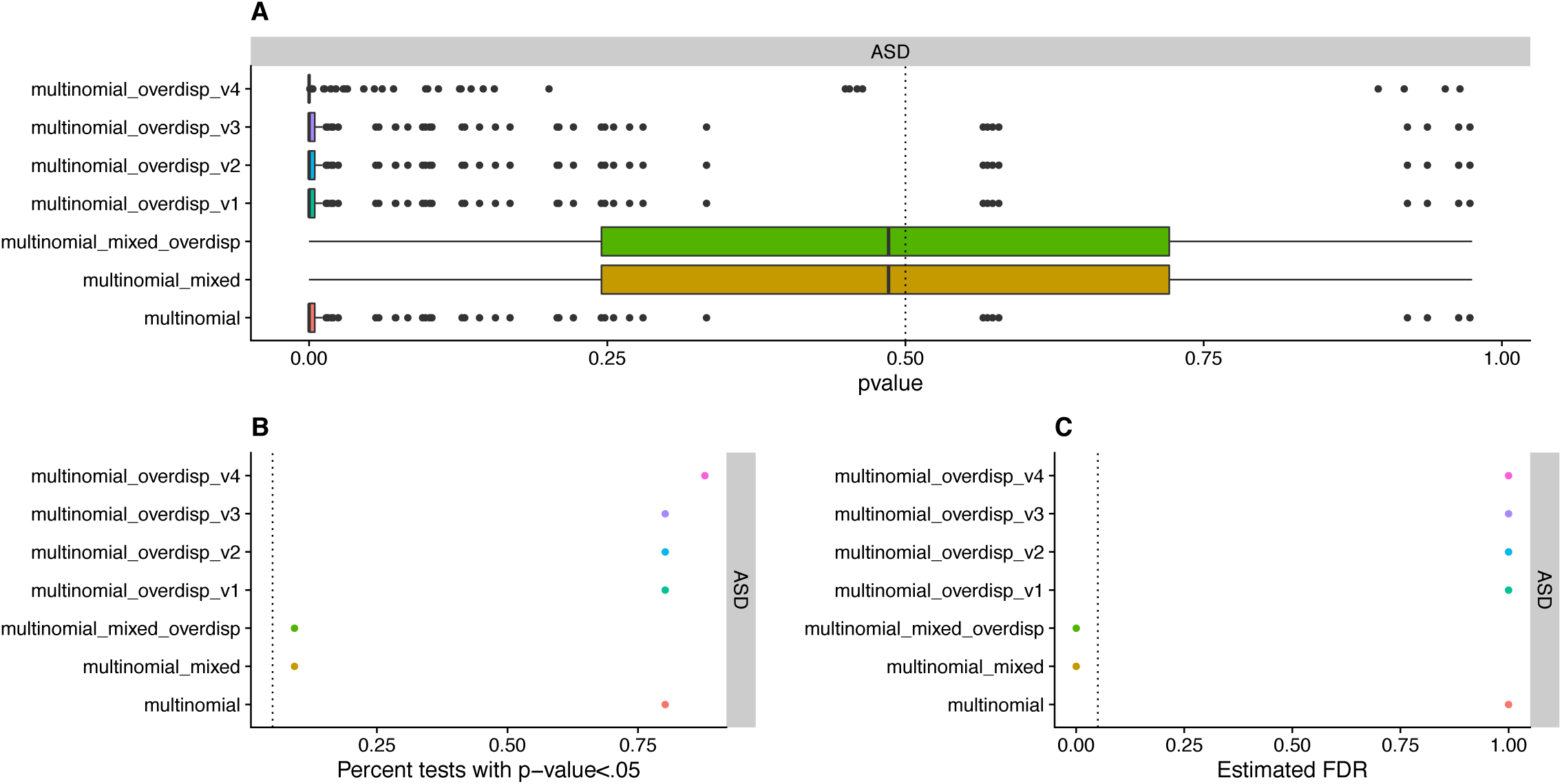
**Overdispersion analysis.** We ran the same analysis as in figures 1 and 2 with the overdispersed multinomial methods. We found the mixed model performed well but the other methods performed poorly with high FPR.

### Effect of test choice on real biological questions

To finish up, we wanted to see how big a difference the choice of method had on various outside datasets where authors aimed to test differences in cell type composition. In particular, we wanted to see if the results extracted were methods specific or not.

To test this we choose a few datasets:

1. Suv Organoid dataset: This dataset consists of organoids, half of which have a Suv420h1 mutation, half do not [18]. There are 3 organoids in each condition.
2. UC dataset: This dataset consists of data from UC patients and healthy controls [5]. Note we only used healthy donor tissue and inflamed UC tissue from the LP, so we expect some differences from their analysis.
3. Aging mouse brain dataset: This data consists of single cell data from mice at 2 different time points [9].

Note that the multinomial based methods failed to run on the UC and Aging mouse data, so the multinomial models were excluded in those analyses. We believe this is likely due to issues with the implementation, not the underlying method, and hope to try to alleviate it in the future.

With these datasets downloaded, it was easy to run all the different methods on them and see (after FDR correction) which cell types were flagged as having significant changes in composition by each method. The results are plotted in Fig 5. In Fig 5a, for each dataset, we have a dot plot of cell type by methods, with dots colored red if that cell type is significant (FDR<.05) with that method, black otherwise. In Fig 5b we also look at the number of significant cell types for each method in each dataset.

**Fig 5:**
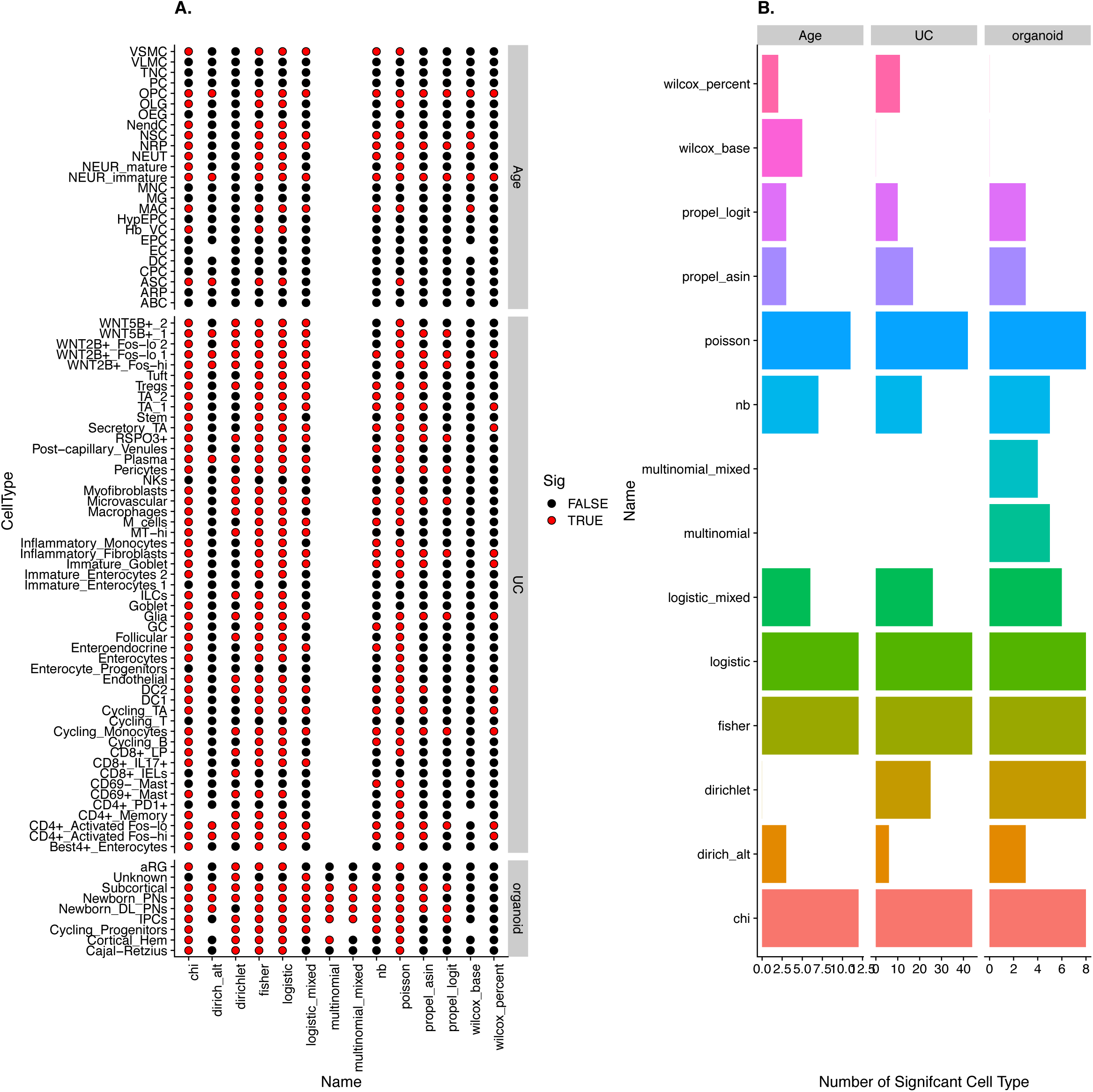
Reanalyzing pervious single cell data results. For each sample, we compared the two conditions with each method (excluding the multinomial based methods in the UC dataset). a) For each dataset, we plot cell type versus method. A black dot indicates a significant results (FDR<.05), a black dot indicates it is not significant, and a white dot indicates it is missing (either due to being the base cell type or the multinomial issue mentioned above). b) For each method in each dataset, a bar whose length is equal to the number of significant cell types. We see large differences between methods in all datasets.

For the Organoid data, the analysis used in the original paper was the logistic_mixed method. Here we see huge differences based off method—wilcox based methods seem to result in no results, while other methods declare almost all cell types as significant. The most interesting result is the dirichlet regression, were the only non-significant cell type is Newborn_DL_PNs, which is one of the top hits with most other methods/one of the ones followed up in the manuscript. We see similar large variation between methods in both the UC (which used the Dirichlet approach in their paper, though a slightly different set up to ours) and age datasets (which used the Wilcox_Percent approach in their paper). It is worth mentioning that, for methods that have both base cell type versions and non-base cell type versions (the wilcox and dirichlet based methods) we see very few significant results in the UC dataset with the base type version compared to the non-basetype version.

We can also look and see which methods found known cell type composition changes, giving us more insight into each methods statistical power in the real world. In the aging dataset, there are three known changes in cell type from prior literature the authors pointed out. Two of them, OPC and Neur_Immature, are found to be significant by every method except dirichlet, while the remaining one, NRP, is found by all of them except dirchlet, dirich_alt, and wilcox_percent. By a similar logic, looking at known cell type changes for UC (mentioned in Smille et al), we see that changes in Tregs are significant by all methods except dirich_alt, the wilcox based ones, and propel_logit, while changes in Mast cells are found only by dirichlet, chi, fisher, logistic, and poisson methods. It is also known that *CD8*^+^*IL-17*^+^ T cells show changes, which is found by the logistic mixed model, chi, logistic, and fisher. Note our results on the UC dataset are slightly different than in the original publications since we used a subset of the original data. This analysis shows that no one method among those with low FPR is consistently able to find known cell type composition changes.

### Robustness and Downsampling Analysis

As a final consideration, we also want to test for robustness to downsampling—how consistent are the results when we downsample the number of samples/cells/etc. One would hope downsampling would not lead to huge changes in the results, and in particular should not lead to many more significant hits. This can also help get at power for lower sample sizes in a more natural way than in our earlier power analysis—assuming the results in the full analysis are correct, we would hope they could be recovered by analysis on the downsampled datasets.

To investigate this, for the Organoid datasets (we left out UC and aging due to the issues with multinomial models), we subsampled 25% of cells at random and reran the analysis from Fig 5. We repeated this 100 times. For each method, among the cell types that where declared significant by that method using the whole dataset we asked what percentage of the time those cells were declared significant in the downsampling analysis. These results are in Fig 6a, with the top plot (pink bars) representing genes that are not significant in the analysis of the whole dataset, the bottom (blue bars) representing those that were significant (note some methods only appear in one or the other since they either had no significant cells types or all cell types significant). We would like the plot on the top (the pink bars) to be close to 0 and the plot on the bottom (the blue bars) to be close to 1. We can see that for almost all methods the pink bars are at/close to 0, while the blue bars range from .5-1.0, with the best performers being dirich_alt, multinomial, and propel_asin. We also did a similar analysis except split the results by cell type, see Fig 6d.

**Fig 6:**
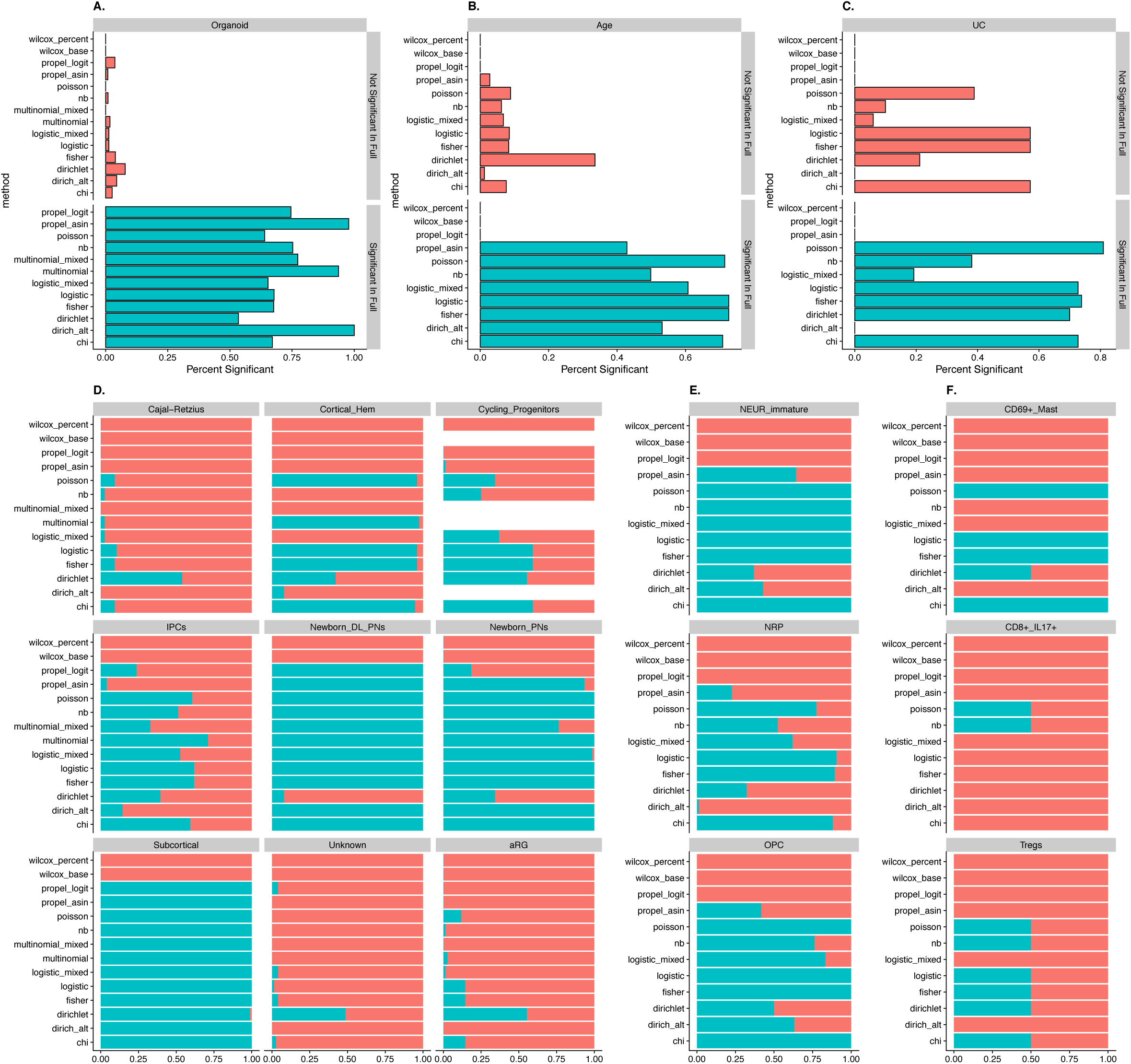
Downsampling analysis a) For the Organoid dataset and each method we split the cell types into those that where declared significant in the entire dataset and those that were not. We then downsampled the cells 100 times and reran the analysis. The pink bars represent the average percent of hits that are not significant in the full analysis that are significant in the downsampled analysis, the blue the percent of ones that are significant in the full analysis that are also significant in the downsampled one. We would like pink bars to be short (so downsampling does not introduce new results), blue bars to be long (so downsampling still allows us to find the results from the full dataset). b) and c) are similar, except for the aged and UC dataset, respectively, and downsampling number of samples instead of number of cells. d-f) For each method and each selected celltype, looks at the percent of downsampling runs where that cell type is significant for that methods, where blue represents signficiant, pink not significant. Here d) is the Organoid dataset with downsampling of cells, e) the aging datasets with downsampling of samples, and f) the UC dataset with downsampling of samples. For e) and f) cell types with known changes are shown.

In addition to the above, we ran a similar analysis with the aging mice dataset and UC dataset, except instead of subsampling cells, we subsampled samples: for each iteration we randomly picked 4 samples from each condition to run the analysis on. Again, we ran this for 100 iterations, though this time we excluded the multinomial based methods due to the issues mentioned previously with them crashing on certain inputs. We do a similar analysis to that in Fig 6a, with Fig 6b being the aging dataset and Fig 6c being the UC dataset. For the aging dataset we see that most methods have pink bars close to 0 as desired, with the exception of dirichlet which is close to .5, while the blue bars are close to .8 mostly for the low performing methods from Fig 1, with most the other methods in the range .4-.6 except for the wilcox based methods and propel_logit which are close to 0. Looking at a similar analysis for the UC dataset, we see the low performing methods from Fig 1 have pink bars near .6, while most the other methods have them near 0 as desired. The results for the blue bars are similar to those for the aging dataset, except propel_asin and dirich_alt are near 0.

It is particularly interesting to look at how often known effects are picked up, since we can be sure they are not false positives and can give us some estimate of power. In the case of both the aging and UC datasets there are 3 cell types with known changes according to the publications, so we look at those to see how often they are detected (a way to test power). The results for aging are in Fig 6E and those for UC are in 6F. Both show mixed results for the sample aware methods, while the methods that are not sample aware (as well as poison) almost always detect these cell types (likely because they declare most cell types significant). It is also interesting to note that the effects in the aging dataset are more often detected than those in the UC dataset. Overall all the high performing methods seem fairly robust, with some exceptions. In the future it might be interesting to build on this analysis to investigate the trade off between the number of cells and samples, and to see if some methods perform better in some circumstances than others.

### Takeaways

The above analysis lends itself to many different interpretations. There is no clear winner, with different methods having different advantages or disadvantages. That being said, there are some clear takeaways:

1. Taking into account sample to sample variability is extremely important—not doing so leads to massive over inflation of p-values. We also see that there is more over dispersion than is accounted for in the Poisson model, suggesting that a different model (NB, etc) should be used.
2. If you are in a situation where choosing a base cell type is acceptable, than doing so alleviates many of the limitations of other methods (this is particularly obvious in the dirichlet case, where the standard parameterization seems to struggle in certain cases), including allowing for dealing with compositionality. Thoughts on how to choose the base cell type are below.
3. If you are unwilling to choose a base cell type, it is not clear that the compositionality aware methods (dirichlet regression with the standard parameterization) are better than the top performing non-compositionality based methods (the propel_asin method) in all cases (such as in the case when one wants to correct for confounders) and can lead to very different results than other methods.
4. Though all methods seem to run in a reasonable time on our data (worse case on the order of a minute), there is a huge variability in the runtime (several orders of magnitude) which may be worth considering in some use cases (when one is using millions of cells, for example).
5. In some situations, numerical issues can lead to methods crashing—we saw this with the multinomial based methods in particular, though is likely to effect most the regression based methods in at least some circumstances. Note that these are not necessarily a limitation of the statistical methods themselves, but instead of the implementations, and we are currently looking into alternative implementations for the multinomial models for those reason. Still, it is worth being aware of these limitations.
6. We see that many of the issues with the Dirichlet regression based approach occur in situations with relatively few biological replicates (3-4 per condition). This suggests the possibility that it may be that the best methods depends on the number of replicates present (though more investigation is required to ensure this).
7. Overall, the data suggests the best methods to use are the Dirichlet based methods, the propel based methods, and the multinomial mixed model, each with its own strengths and weaknesses. Probably the most consistent performers were propel_asin and dirich_alt, though it is unclear if they are the best choices in all situations (in particular, due to lack of explicit compositionality, it is unclear what happens to propel_asin in more extreme version of the power test).

We put the results into one figure, Figure 7, with the caveat that the colors are based off our interpretation of the results and different people might disagree. Green represents high performance, yellow represents mixed, and red poor performance. Grey represents performance that could not be fairly measured (mostly due to issues getting the method to run).

**Fig 7:**
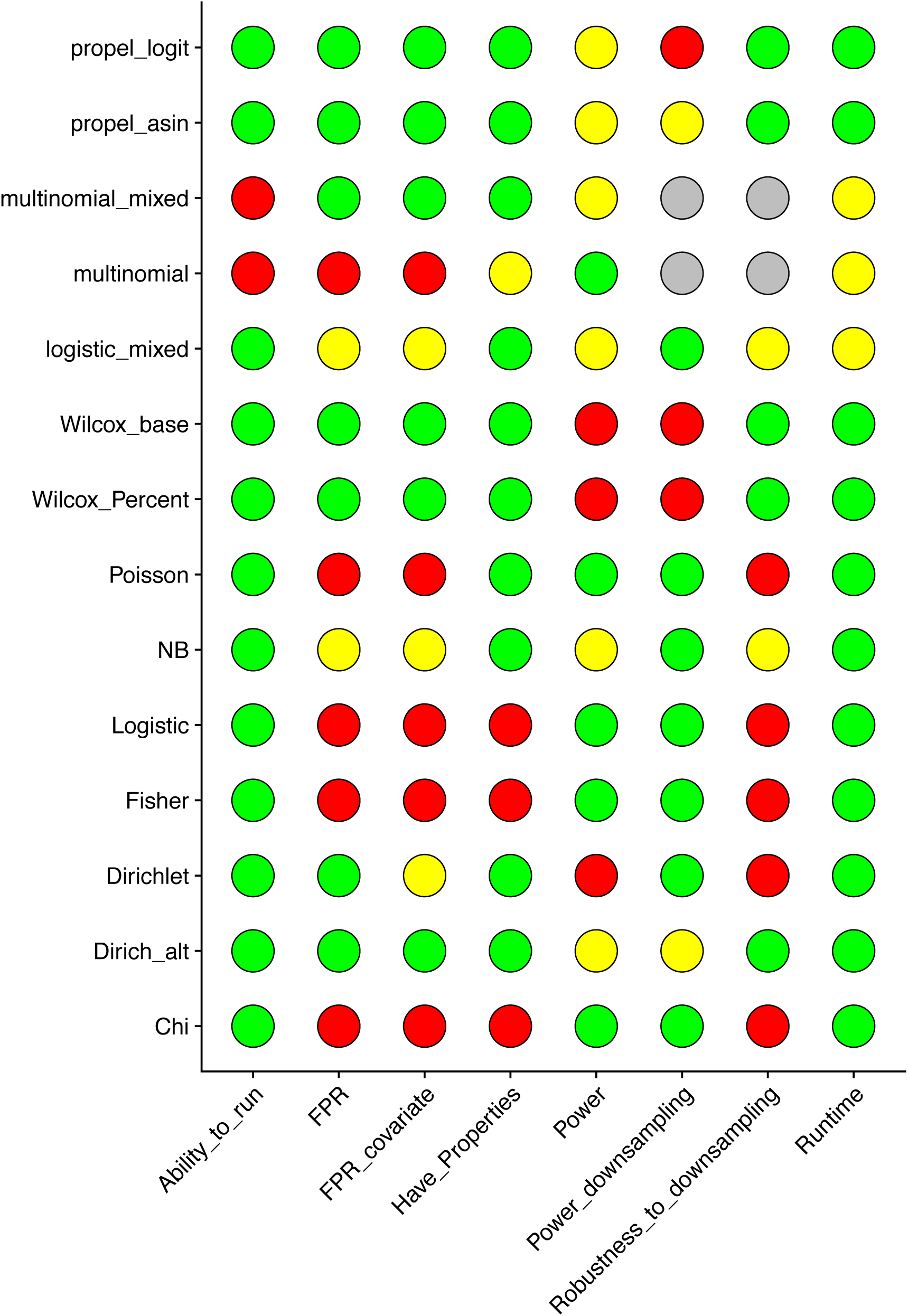
Overall performance. Here green represents high performers, yellow medium/mixed performers, and red poor performers. Grey represents cases where we could not run the method to test.

If one decides to use a method with a base cell type, this raises the question of how to choose that cell type. If there is an obvious biological choice for base that is of course the best approach (for example, if one is interested in changes in glia choosing excitatory neurons as the base might make sense). Another obvious option is choosing the cell type with the most cells (can give you more power). One could also try to do a model selection like approach to choose the base cell type—choose the one that gives the model with the smallest regression coefficients (aka smallest L1 or L2 values), for example, or use a regularization based approach (lasso, etc, something we did not explore here). Finally, one could try using each cell type as the base, and instead of looking for which cell types differ and which don’t focus on which cell types change relative to one another—in many ways this might be the most reliable/interpretable method.

It is worth noting this is far from a complete comparison—for example it focuses exclusively on p-values and ignores effect size estimates, much of the analysis is based on datasets with a fairly small number of samples (the ASD data set in particular), etc. More work is left to be done in the future!

### Method

All methods were implemented in R. The R packages used for each test are listed in the **Description of methods** section. Code is also in the process of being made available on github for more details (see the Code Availability section). FDR correction was performed with the “fdr” option in p.adjust.

For each analysis data was downloaded from online data sources and processed to create tables containing the meta data for each cell. In particular, the meta data for each dataset contained cell type annotation and sample of origin. For samples with different conditions (Aging, Organoid, and UC datasets) condition information was recorded as a column as well.

For permutation testing, each sample was randomly assigned to either be a knockout (ko) or a wild type (wt) such that half were assigned to each condition. For analysis of covariates, samples were randomly assigned to have the covariate (assigned 1) or not (assigned 0) such that half were assigned to each. For power analysis, after the condition was assigned, we randomly sampled Excitatory neurons in ko samples to be removed (either with probability .05 or .5). For the downsampling analysis, either cells were selected to be included randomly with probability .25 (for the Organoid data) or 8 samples were selected at random (for the other datasets).

For the Propeller method we modified their code to take in data in the format we are using.

For each dataset generated as above each cell type composition test was run (details on their implementation are in **Description of methods** section). The package tictoc was used to get information about runtime.

All visualization was performed with the ggplot2, cowplot, and patchwork packages.

## Acknowledgements

Thanks to Joshua Levin for giving me the time to work on this, and for Joshua Levin, Kwanho Kim, Amanda Kedaigle, Adam Haber, and others for helpful discussions/insight.

## Data Availability

All data used was from previously published studies. Location data was downloaded from:

***Immune data:*** https://singlecell.broadinstitute.org/single_cell/study/SCP548

***Organoid data:*** https://singlecell.broadinstitute.org/single_cell/study/SCP1129

***UC data:*** https://singlecell.broadinstitute.org/single_cell/study/SCP259

***Age data:*** https://singlecell.broadinstitute.org/single_cell/study/SCP263

***ASD data:*** https://cells.ucsc.edu/?ds=autism

## Code Availability

Code (not up to data/not well documented, work in progress) available at: https://github.com/seanken/CellTypeComp/

## Notes

### Competing Interest Statement

The authors have declared no competing interest.

